# The cortical hubs related to recovery of consciousness

**DOI:** 10.1101/2021.04.25.441310

**Authors:** Hang Wu, Zengxin Qi, Jun Zhang, Changwei Wu, Xuehai Wu, Zirui Huang, Di Zang, Stuart Fogel, Sean Tanabe, Anthony G. Hudetz, Georg Northoff, Ying Mao, Pengmin Qin

## Abstract

**Background and Objectives:** The neural mechanism that enables the recovery of consciousness in patients with unresponsive wakefulness syndrome (UWS) remains unclear. The aim of the current study is to characterize the cortical hub regions related to the recovery of consciousness in patients with UWS.

**Methods:** Voxel-wise degree centrality analysis was adopted to identify the cortical hubs related to the recovery of consciousness, for which a total of 27 UWS patients were used, including 13 patients who emerged from UWS (UWS-E), and 14 patients who remained in UWS (UWS-R) at least three months after the experiment performance. Furthermore, other recoverable unconscious states including three independent deep sleep datasets (n = 12, 9, 9 respectively) and three independent anesthesia datasets (n = 27, 14, 6 respectively) were adopted as validation groups. Spatial similarity of the hub characteristic with the validation groups between the UWS-E and UWS-R was compared using the dice coefficient. Finally, with the cortical regions persistently shown as hubs across UWS-E and validation states, functional connectivity analysis was further performed to explore the connectivity patterns underlying the recovery of consciousness.

**Results:** Four cortical hubs were identified with significantly higher degree centrality for UWS-E than UWS-R, including the anterior precuneus, left inferior parietal lobule, left inferior frontal gyrus, and left middle frontal gyrus, of which the degree centrality value also positively correlated with the patients’ Glasgow Outcome Scale (GOS) score. Furthermore, the anterior precuneus was found to show significantly higher similarity of hub characteristics as well as functional connectivity pattern between UWS-E and validation groups, compared with UWS-R.

**Discussion:** The results suggest that the recovery of consciousness may be relevant to the integrity of cortical hubs, especially the anterior precuneus. The identified cortical hub regions could serve as potential targets for noninvasive stimulation aimed at promoting the patients’ consciousness recovery.

## Introduction

Disorders of consciousness (DOC) are usually due to severe brain damage, which includes unresponsive wakefulness syndrome (UWS) and minimally conscious state (MCS).^1^ Of all UWS patients, some could eventually regain consciousness, and some don’t.^2^ The identification of robust prognostic markers for the recovery of consciousness in UWS has significant values in the fields of clinical and basic researches.^3^ However, researches done on UWS is challenging, due to their lack of behavioral responsiveness.^4^ With resting-state fMRI (rs-fMRI), the residual awareness^5–7^ could be detected within patients with UWS by measuring the brain connection within large-scale brain networks, which could be useful to predict their recovery of consciousness.^8,9^

Graph theory is one of the methods that has been applied in studying the connection within or between brain networks for patients with DOC,^10,11^ due to the fact that functional integration could play a crucial role in supporting consciousness.^12–14^ In the graph theoretical analysis, cortical hub regions were defined as nodes with much high functional connectivity with other nodes in the brain,^15^ which could promote the functional integration across various components of the network.^16^ Degree centrality is a common metric developed based on graph theory,^17^ in which a higher degree centrality value reflects a larger number of connections between a node (voxel) and the rest of the nodes in the brain, which could facilitate frequent information exchange for healthy brain functioning,^18^ and a cluster of nodes with high degree centrality values is considered a hub. Previous studies using this method have found that cortical hubs were largely reorganized in DOC patients compared to fully conscious volunteers.^10,11^ Moreover, previous studies have showed that cortical hubs play a key role in supporting consciousness. For instance, some studies have found some similarity in hub distributions among DOC and other recoverable unconscious states, such as deep sleep and anesthesia.^6,14^ Based on these findings, it is possible that cortical hubs could also play an important role in the recovery of consciousness, which, however, remains uninvestigated. Following this idea, by comparing the difference and similarity in the distribution of cortical hubs across recoverable and unrecoverable UWS patients as well as other recoverable unconscious states (e.g., deep sleep and anesthesia states), it should be a reasonable way to uncover the cortical hubs supporting the recovery of consciousness.

In the current studies, we aim to identify the cortical hubs that could support the recovery of consciousness using two subgroups of UWS patients: patients who emerged from UWS (UWS-E) and patients who remained in UWS (UWS-R) at least three months after the experiment performance. Two recoverable unconscious states were used as validation groups (including three independent groups of deep sleep and three independent groups of anesthesia state), to compare their similarity of the spatial distribution of hubs with UWS patients. Specifically, we first conducted the degree centrality analysis on the two subgroups of UWS patients. After obtaining hub regions that play a crucial role in differentiating the two groups, which were defined as ROIs, the dice coefficient was then used to assess the similarity between the spatial distribution of hubs in UWS-E and validation groups (deep sleep and anesthesia states), as well as UWS-R and validation groups within these ROIs, respectively. Finally, the common cortical hubs across UWS-E, deep sleep, and anesthesia groups were used as seeds, and seed-based functional connectivity was calculated to explore the corresponding brain network related to the recovery of consciousness.

## Methods

### Participants and imaging acquisitions: UWS patients

Thirty-one patients were recruited from Huashan Hospital affiliated to Fudan University (Shanghai, China). Due to excessive head motion, four patients were excluded from subsequent analysis, leaving twenty-seven patients (male/female: 20/7; age range: 17 - 67 years) with structurally well-preserved brain images, which were carefully chosen by author XW and checked by author PQ. Before fMRI scanning, patients with UWS were assessed using Coma Recovery Scale-Revised (CRS-R) scale.^19^ The patients were assessed again at least 3 months after the scanning using the Glasgow Outcome Scale (GOS). Patients with a GOS score ≥ 3 were regarded as emerged from UWS (UWS-E, male/female: 9/4; age range: 17 - 52 years), and patients with a GOS score < 3 were regarded as remained in UWS (UWS-R, male/female: 11/3; age range: 26 - 67 years). No significant difference was found between UWS-E and UWS-R in terms of sex (p = 0.58), age (p = 0.07), days after insult (p = 0.73), and CRS-R score at the time of scan (p = 0.98) (See Table 1 for detailed demographic and clinical information).

**Table 1.** Demographic and clinical information for unresponsive wakefulness syndrome patients

| Patient number | Sex/Age | Cause | Time of fMRI<br>(days after insult) | CRS-R | Diagnosis | GOS |
| --- | --- | --- | --- | --- | --- | --- |
| p1 | F/35 | TBI | 21 | 5 | UWS-E | 4 |
| p2 | F/24 | TBI | 1 | 3 | UWS-E | 4 |
| p3 | M/46 | SIH | 18 | 2 | UWS-E | 4 |
| p4 | F/43 | SIH | 162 | 6 | UWS-E | 3 |
| p5 | M/52 | TBI | 210 | 6 | UWS-E | 3 |
| p6 | M/38 | TBI | 81 | 3 | UWS-E | 3 |
| p7 | M/46 | TBI | 25 | 2 | UWS-E | 3 |
| p8 | M/44 | TBI | 21 | 3 | UWS-E | 4 |
| p9 | F/39 | TBI | 31 | 6 | UWS-E | 3 |
| p10 | M/45 | TBI | 12 | 6 | UWS-E | 4 |
| p11 | M/46 | TBI | 168 | 1 | UWS-E | 3 |
| p12 | M/46 | TBI | 162 | 5 | UWS-E | 3 |
| p13 | M/17 | Anoxic | 493 | 7 | UWS-E | 3 |
| p14 | M/50 | TBI | 137 | 3 | UWS-R | 2 |
| p15 | M/30 | TBI | 259 | 8 | UWS-R | 2 |
| p16 | F/57 | TBI | 15 | 3 | UWS-R | 2 |
| p17 | M/60 | SIH | 109 | 6 | UWS-R | 2 |
| p18 | M/26 | TBI | 143 | 6 | UWS-R | 2 |
| p19 | M/47 | SIH | 315 | 6 | UWS-R | 2 |
| p20 | M/46 | Anoxic | 23 | 3 | UWS-R | 2 |
| p21 | M/51 | TBI | 100 | 4 | UWS-R | 2 |
| p22 | M/53 | TBI | 30 | 3 | UWS-R | 2 |
| p23 | F/67 | TBI | 10 | 3 | UWS-R | 2 |
| p24 | M/57 | SIH | 33 | 2 | UWS-R | 2 |
| p25 | F/52 | SIH | 51 | 4 | UWS-R | 2 |
| p26 | M/39 | SIH | 221 | 4 | UWS-R | 2 |
| p27 | M/35 | SIH | 300 | 4 | UWS-R | 2 |

Note: UWS = unresponsive wakefulness syndrome; UWS-E = patients who emerged from UWS at least three months after the experiment performance; UWS-R = patients who remained in UWS at least three months after the experiment performance; CRS-R = Coma Recovery Scale-Revised; GOS = Glasgow Outcome Scale; SIH = spontaneous intracerebral hemorrhage; TBI = traumatic brain injury.

In this dataset, the MR images were acquired using a Siemens 3 Tesla scanner. Functional images were acquired using a T2*-weighted EPI sequence (TR/TE/θ = 2000ms/35ms/90°, FOV = 256 × 256 mm, matrix = 64 × 64, 33 slices with 4-mm thickness, gap = 0 mm, 200 scans). For functional image registration and localization, a high-resolution T1-weighted anatomical image was also acquired for each participant.

### Participants and imaging acquisitions

In order to validate the results obtained from UWS group, additional unconscious states were collected, including the anesthesia and deep sleep state. Specifically, for repeatability validation, three independent datasets of deep sleep and three independent datasets of anesthesia were included.

### Sleep group 1

This dataset was analyzed out of a different purpose in a previously published study.^20^ Twelve healthy men (ages 20 to 27) were included, who slept for 7-8 hours each night and had a consistent bed/wake time for four days. For the sleep protocol, simultaneous EEG–fMRI was recorded. The sleep protocol was conducted at midnight and the participants were told to try to fall asleep as soon as the EPI scan began. A licensed sleep technician at Kaohsiung Medical University Hospital visually rated sleep stages for every 30-second interval, according to the American Academy of Sleep Medicine’s current criteria (AASM).

The MR images were acquired on a 3 Tesla Siemens Tim-Trio scanner. Functional images were acquired using a T2*-weighted EPI sequence (TR/TE/θ = 2500ms/30ms/80°, FOV = 220 × 220 mm, matrix = 64 × 64, 35 slices with 3.4 mm thickness). A high-resolution T1-weighted anatomical image was acquired on all participants for functional image registration and localization. More details on the EEG and fMRI data acquisition parameters and preprocessing can be found in the previous analysis.^20^

### Sleep group 2

The dataset was analyzed out of a different purpose in a previously published study^21^ using simultaneous EEG–fMRI recordings. In the current study, nine healthy participants with a persistent N3 sleep stage (male/female: 3/6; age range: 18 - 33 years) were included. The sleep stages were scored in the same way as Sleep group 1. The MR images were acquired on a 3 Tesla Siemens Magnetom Prisma scanner. Functional images were acquired using a T2*-weighted EPI sequence (TR/TE/θ = 2160ms/30ms/90°, FOV = 220 × 220 mm, matrix= 64 × 64 × 40, 40 slices with 3 mm thickness). For image registration and localization, a high-resolution T1-weighted anatomical image was acquired for each participant. More details about the EEG and fMRI data acquisition parameters and data preprocessing can be found in the previous analysis.^21^

### Sleep group 3

The dataset was analyzed out of a different purpose in a previously published study^22^ using simultaneous EEG–fMRI recordings. Nine healthy participants (male/female: 5/4; age range: 22 - 35 years) with a persistent N3 sleep stage were included. The sleep stages were scored in the same way as Sleep group 1 and Sleep group 2. MR images were acquired on a 3 Tesla Siemens Tim-Trio scanner. Functional images were acquired using a T2*-weighted EPI sequence (TR/TE/θ = 2160ms/30ms/90°, FOV = 220mm, matrix size = 64 × 64, slice thickness = 3 mm, 40 slices). A high-resolution T1-weighted anatomical image was acquired for each participant for registration and localization. More details on the EEG and fMRI data acquisition parameters and preprocessing can be found in the previous analysis.^22^

### Anesthesia group 1

The dataset was analyzed out of a difference purpose in previously published studies before.^23,24^ Twenty-seven participants received intravenous propofol anesthesia (male/female: 13/14; age range: 27 - 64 years). The elective transsphenoidal approach for resection of pituitary microadenoma had been used on all of participants.

A plasma concentration of 3.0-5.0 g/ml was achieved using a target-controlled infusion (TCI) based on the Marsh model. Unconsciousness was reliably induced using TCI propofol at a stable effect-site concentration (4.0 g/ml) for each participant. A participant was considered unconscious (Ramsay 5–6) if he/she did not respond to verbal commands such as “squeeze my hand” during anesthesia. Furthermore, in the post-operative follow-up, no participant reported explicit recollection. Therefore, all participants were deemed unconscious during anesthesia. Participants were given intermittent positive pressure ventilation during the anesthetic state, with a tidal volume of 8-10 ml/kg, a respiratory rate of 10-12 beats per minute, and a PetCO2 (end tidal partial pressure of CO2) of 35-37 mmHg. Throughout the study, two certified anesthesiologists were present to ensure that resuscitation equipment was always accessible. See the previous study for more details of the anesthesia protocols.^23,24^

A Siemens 3T scanner (Siemens, Erlangen, Germany) was used to obtain functional photographs of the whole brain using a T2*-weighted EPI series (TR/TE/θ = 2000ms/30ms/90°, FOV = 192 mm, matrix size = 64 × 64, 25 slices with 6-mm thickness, gap = 0 mm, 240 scans). A high-resolution T1-weighted anatomical image was acquired for each participant for functional image registration and localization. The collection of EPI data was carried out in both in an awake state prior to anesthesia and in an anesthetic state.

### Anesthesia group 2

An independent dataset during propofol-induced deep sedation was included, which was analyzed out of a different purpose in previously published studies.^23,24^ One participant were excluded due to excessive head motion, leaving fourteen participants (male/female: 8/6; age range: 19-34 years) in the current study.

The levels of behavioral responsiveness were measured using the OAAS (observer’s assessment of alertness/sedation). Participants responded readily to verbal commands during baseline conscious and recovery conditions (OAAS score, 5). Participants responded to verbal commands with a sluggishness when under light sedation (OAAS score, 4). During deep sedation, participants showed no response to verbal instruction (OAAS score, 2 and 1). The resulting target plasma concentrations differ among participants (light sedation, 0.98 ± 0.18 μg/ml; deep sedation, 1.88 ± 0.24 μg/ml) due to the variability of individual anesthetic susceptibility. During anesthesia, the plasma concentration of propofol was maintained in equilibrium by constantly changing the infusion rate to maintain the balance between accumulation and drug elimination. The infusion rate was manually monitored and driven by the results of a computer simulation based on the pharmacokinetic model of propofol that was designed for target-controlled drug infusion (STANPUMP). During the procedure, the electrocardiogram, noninvasive blood pressure cuff, pulse oximetry, and end-tidal carbon dioxide gas monitoring were all performed according to standard American Society of Anesthesiologists (ASA) protocols. A nasal cannula was used to administer supplemental oxygen as a preventative measure. For more details on the anesthesia procedures, see the previous studies.^23,24^

Functional images of the whole brain were acquired using a 3T Signa GE 750 scanner (GE Healthcare, Waukesha, WI, USA) that used T2*-weighted EPI sequence (TR/TE/θ = 2000ms/25ms/77°, FOV = 224 mm, matrix size = 64 × 64, 41 slices with 3.5 mm thickness, 450 scans). A high-resolution T1-weighted anatomical image was acquired for each participant. The data acquisition of EPI was carried out in a baseline-conscious state, light and deep sedation and recovery.

### Anesthesia group 3

Six participants who received inhalational sevoflurane anesthesia (male/female: 3/3; age range: 26 - 62 years) were included. The elective transsphenoidal approach for resection of pituitary microadenoma was used on all of participants.

We administered 8% sevoflurane in 100% oxygen with a fresh gas flow of 6 L/min to the sevoflurane group, along with remifentanil (1.0 μg/kg) and succinylcholine (1.0 mg/kg). This was maintained with 2.6% (1.3 MAC) ETsevo in 100% oxygen, and a fresh gas flow of 2.0 L/min. Sevoflurane concentration effectively preserved a lack of consciousness in participants identified as ASA physical status I or II. The rest of the anesthesia procedures and the acquisition parameters of fMRI were the same as the Anesthesia group 1.

### Preprocessing

fMRI data was preprocessed using AFNI (Analysis of Functional NeuroImages), including: discarding the first two volumes; slice timing correction; head motion correction; co-registration of functional and structural images; non-linear transformation to Montreal Neurological Institute (MNI) template; a volume and previous volume were labeled as 0 (1, otherwise) if its derivative values of six-dimensional motion parameters have a Euclidean Norm (square root of the sum squares) above 0.5;_25_ band-pass filter was applied between 0.01 and 0.1 Hz; various uninterested components were removed via linear regression, including time series of head motion and its temporal derivative, mean time series from white matter and ventricle, and binarized censored time series (output data included zero values at censored time points); spatial smoothing was applied using a Gaussian filter with a full width at half maximum of 6 mm.

### Degree centrality analysis

According to the graph theory, the number of edges connecting a node with other nodes reflects the importance of a node, which is defined as its degree centrality.^26^ Based on this definition, each voxel was considered a node, and the functional connection (Pearson correlation of BOLD time series) between the voxel (i.e. node) and any other voxel could be considered an edge. In this study, a binary mask was created using the Human Brainnetome Atlas which includes the grey matter and subcortical regions,^27^ and then a voxel-wise degree centrality map for each participant was generated using the 3dDegreeCentrality program in AFNI. Specifically, a voxel-wise correlation matrix was obtained for each participant by calculating the correlation coefficient between any pair of voxels. This matrix was then binarized by applying a threshold at r > 0.3 for each participant, and the degree centrality was calculated as the sum of connections for every voxel. We also calculated degree centrality maps at r > 0.2 and r > 0.4 respectively, to validate the effect of different thresholds.

To identify cortical regions related to the recovery of consciousness, the degree centrality maps were compared between UWS-E and UWS-R subgroup. Specifically, independent samples t-test was performed using 3dttest++ program in AFNI with a cluster-level significance threshold of p < 0.05 after FWE correction (p < 0.005 uncorrected, cluster size > 91 voxels, 3dClusterSim, AFNI). In this step, patients with mean degree centrality values beyond the range (mean ± 2SD) for each group were considered as an outlier and excluded from further analysis, which resulted in the exclusion of one UWS-E patient and one UWS-R patient. After the comparison, regions with a significant difference of degree centrality values were obtained, which was used as ROIs (regions of interest) for further analyses.

Furthermore, to verify the relation between the degree centrality values within the ROIs and the patients’ GOS scores, partial Spearman correlation was calculated, in which the global mean of degree centrality value was treated as a confounding variable. All p values of the spearman correlation coefficient were FDR corrected.

### Comparison of cortical hub distribution between UWS-E and UWS-R

To see whether the same distribution of hubs was maintained in UWS-E and UWS-R, ROIs obtained from the degree centrality analysis were used as masks, and the number of overlapped voxels between these ROIs and the distribution of hubs for each individual participant was calculated. For that we first normalized the degree centrality scores into Z scores with zero mean and unit variance for each individual participant, turning a degree centrality map into a Z score map, where cortical hubs were defined as voxels with z values over 1.^18,26^ Next, we calculated the number of overlapped voxels between the ROIs and the hubs found in each individual map for the UWS-E and UWS-R patients respectively. Finally, for each ROI, the numbers of these overlapped voxels for each individual were compared between the two UWS subgroups using independent sample t-test. All p values above were FDR corrected.

### Validation of the UWS result with sleep and anesthesia states

First, the spatial distribution of cortical hubs was computed for all six validation groups (three independent deep sleep datasets and three independent anesthesia datasets). Specifically, to minimize individual differences, the degree centrality map for each participant was first normalized, dividing voxel degree by the global mean degree. Then, normalized degree centrality maps within each group were averaged and transformed to a Z score map. The cortical hubs for each group were defined as voxels with z values over 1, thus creating a binarized map of cortical hubs for each group. Then, the dice coefficient^28^ was used to measure the spatial similarity of the binarized cortical hub maps between the two groups, which measured two times the number of overlapped voxels divided by the total number of voxels in both binary maps, and has a value that ranges from 0 to 1, in which 1 denotes perfect overlapped voxels and 0 denotes no overlapped voxels.^29^ Specifically, within the range of each ROI, six pairwise combinations of dice coefficient between UWS-E and validation groups, and six pairwise combinations of dice coefficient between UWS-R and validation groups were computed respectively. Then Mann–Whitney U test was used to compare the difference of dice coefficients. All p values above were FDR corrected.

In the last step, we used the common hub regions identified in the previous steps as seeds, which showed high similarity of hub characteristics among UWS-E, deep sleep and anesthesia, to calculate the functional connectivity for all groups, thus obtaining the connectivity patterns under various unconscious states. For each group, the connectivity pattern was obtained by applying a threshold with a Z value at 0.3095 (r = 0.3). Then dice coefficient was used again to measure the spatial similarity of the connectivity patterns between UWS-E and validation groups, as well as between UWS-R and validation groups. A permutation test with 10000 replicates was performed to compare the difference of dice coefficients. In addition, to see the connectivity pattern under conscious state, fMRI data of eighty-eight healthy volunteers in their conscious state was obtained (in which 77 came from the participants used for the unconscious state in the validation analysis), and the connectivity pattern was also calculated using the same seeds in the last step to serve as a reference template.

### Standard Protocol Approvals, Registration, and Patient Consent

For UWS group, informed written consent was obtained from legal representatives of the patients, and the study was approved by the Ethics Committee of Huashan Hospital. For Sleep group1, informed consent was obtained from all participants, and ethical approval was obtained from the Institutional Review Board of National Yang-Ming University. For Sleep group 2, informed consent was obtained from all participants and ethical approval was obtained from the Health Science research ethics board of Western University. For Sleep group 3, informed consent was obtained from all participants and ethical approval was obtained from the Research Ethics Board at the Institut universitaire de gériatrie de Montréal (IUGM). For Anesthesia group1 and Anesthesia group 3, informed written consent was obtained from each participant, and the study was approved by the Ethics Committee of Shanghai Huashan Hospital, Fudan University, Shanghai, China. For Anesthesia group 2, informed written consent was obtained from each participant and the study was approved by the Ethics Committee of Medical College of Wisconsin (MCW).

## Data availability

All data needed to evaluate the conclusions in the paper are present in the paper. Additional data related to this paper may be requested from the corresponding author.

## Results

### Cortical regions related to the recovery of consciousness in patients with UWS

Firstly, four cortical regions showed higher degree centrality values in the UWS-E > UWS-R contrast, including the anterior precuneus (PCun), left inferior parietal lobule (LIPL), left inferior frontal gyrus (LIFG), and left middle frontal gyrus (LMFG) (FWE corrected, p < 0.005, cluster size > 91 voxels) (figure 1A-D, left column). In addition, this result was consistent with ones of degree centrality maps with r > 0.2 or r > 0.4.

**Figure 1.**
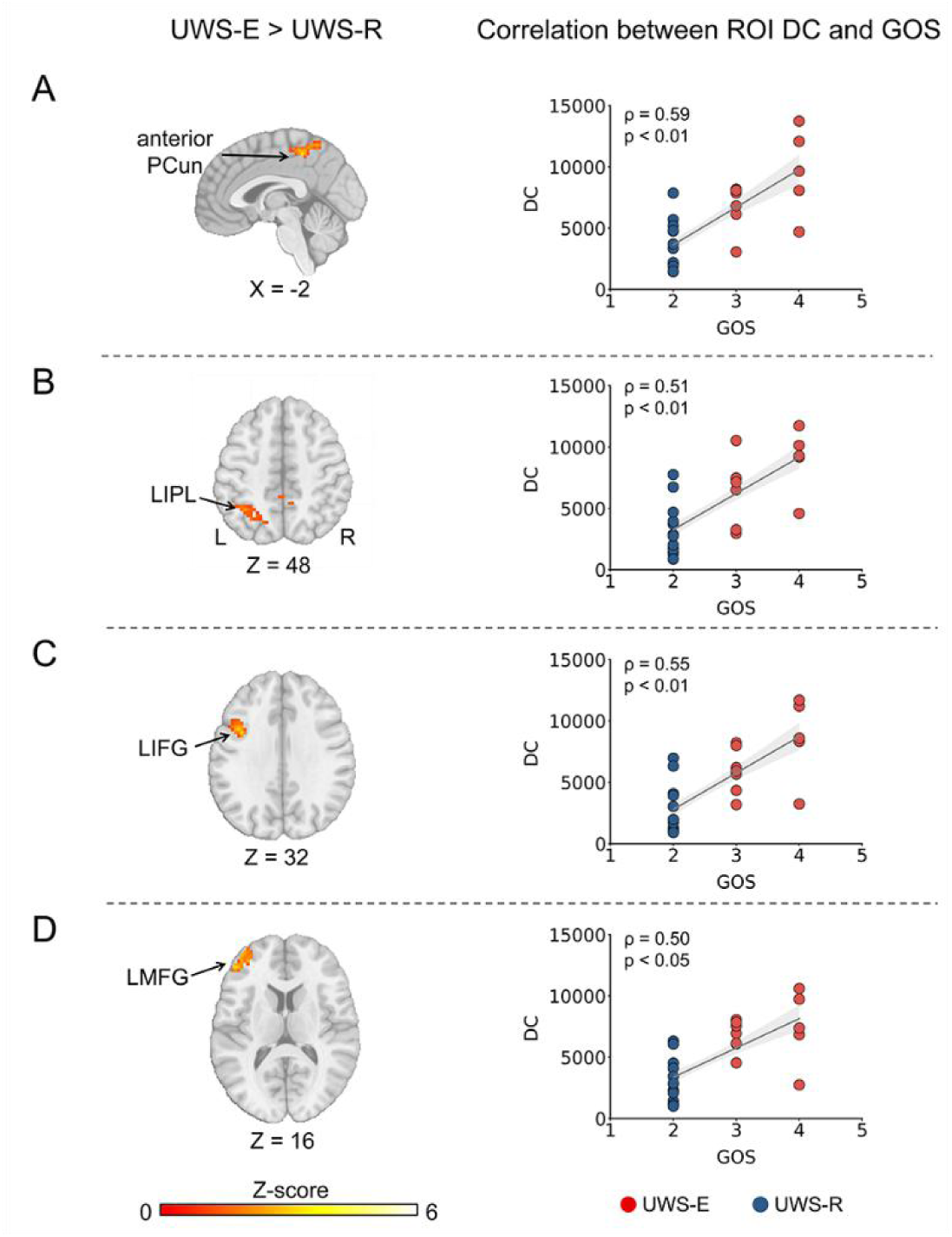
Cortical regions with increased degree centrality in UWS-E than UWS-R (left panel) and their correlation with clinical outcomes (right panel). Four cortical regions obtained including the anterior PCun (A), LIPL (B), LIFG (C), and LMFG (D). PCun = precuneus; LIPL = left inferior parietal lobule; LIFG = left inferior frontal gyrus; LMFG = left middle frontal gyrus; UWS = unresponsive wakefulness syndrome; UWS-E = patients who emerged from UWS at least three months after the experiment performance; UWS-R = patients who remained in UWS at least three months after the experiment performance; DC = degree centrality; GOS = Glasgow Outcome Scale.

Next, partial spearman correlation between the mean degree centrality and GOS scores showed significant positive correlations in the anterior PCun (ρ = 0.59, p < 0.01, FDR corrected), LIPL (ρ = 0.51, p < 0.01, FDR corrected), LIFG (ρ = 0.55, p < 0.01, FDR corrected), and LMFG (ρ = 0.50, p <0.05, FDR corrected) (figure 1A-D, right column).

### Comparison of cortical hub distribution between UWS-E and UWS-R

As shown in figure 2, the spatial distribution of cortical hubs at a group level (blue area) and the contour of all four cortical regions obtained in the last step (red solid line) were visually displayed for the UWS-E (figure 2A-D, left column) and UWS-R (figure 2A-D, middle column). In addition, direct comparison between the two subgroups showed significantly more voxels for UWS-E which overlapped with the four cortical regions identified in the last step (figure 2A-D, right column).

**Figure 2.**
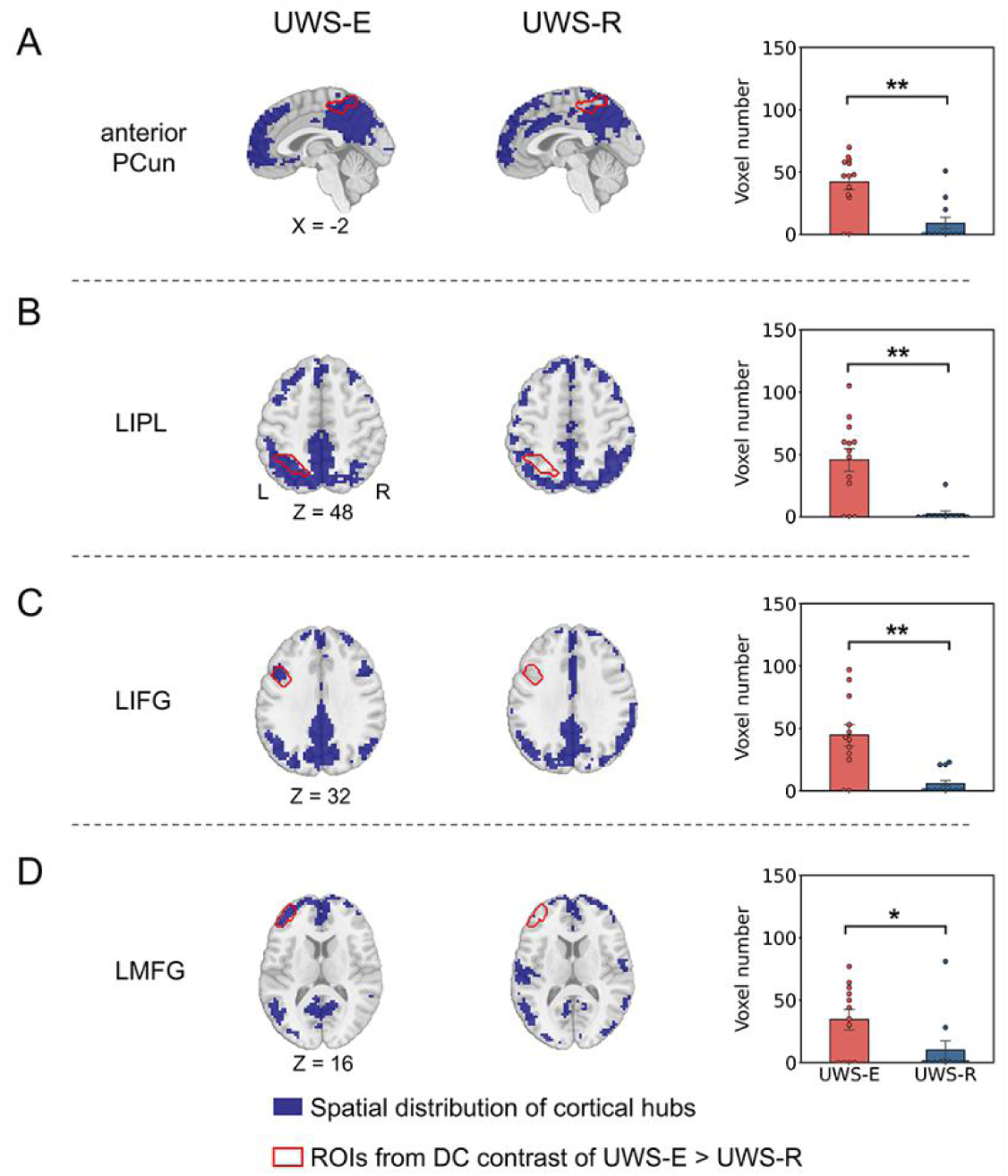
Comparison of hub distribution in UWS patients. For anterior PCun (A), LIPL (B), LIFG (C) and LMFG (D), the distribution of cortical hubs on a group level (blue area) and the contour of cortical regions obtained from the last step (red solid line) was displayed, as well as the number of overlapped voxels between each ROI and the distribution of hubs for each individual participant was calculated and compared between UWS-E and UWS-R. PCun = precuneus; LIPL = left inferior parietal lobule; LIFG = left inferior frontal gyrus; LMFG = left middle frontal gyrus; UWS = unresponsive wakefulness syndrome; UWS-E = patients who emerged from UWS at least three months after the experiment performance; UWS-R = patients who remained in UWS at least three months after the experiment performance; DC = degree centrality; ** indicates p < 0.01 FDR corrected; * indicates p < 0.05 FDR corrected.

### Similarity of cortical hub distribution among UWS-E, deep sleep and anesthesia states

As shown in figure 3, the group-level spatial distribution of cortical hubs (blur area) and all four cortical regions obtained in the first step (red solid line) were visually displayed for all six validation groups. As shown in figure 4, group-level comparison of the dice coefficient showed significantly more similarity with all six validation groups for UWS-E than UWS-R in the anterior PCun (p < 0.05, FDR corrected). In addition, this result was consistent with degree centrality maps with r > 0.2 or r > 0.4.

**Figure 3.**
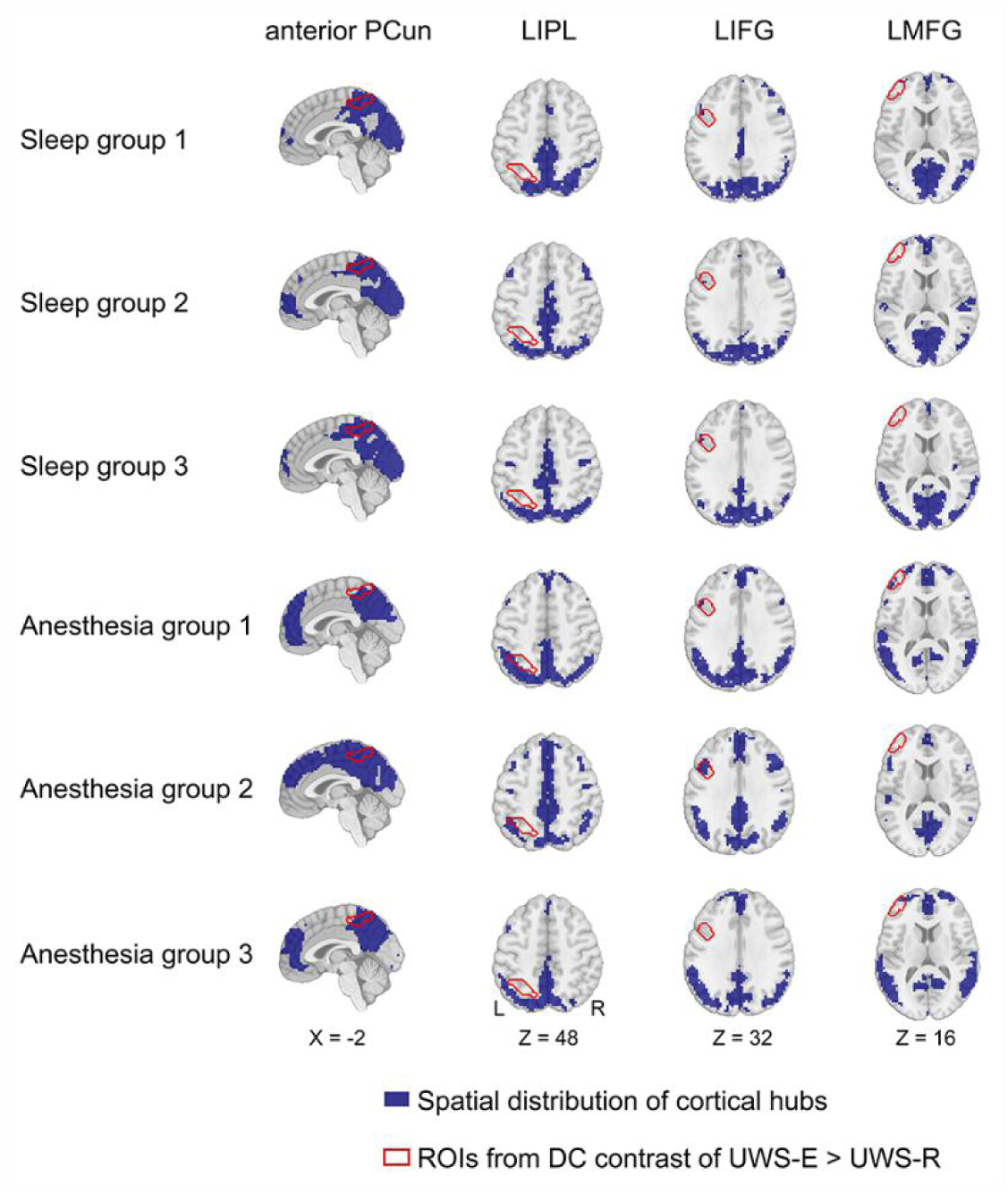
Spatial distribution of cortical hubs for all validation groups. For each group, the spatial map of cortical hubs (blue area) and the contour of the four hubs obtained in the first step (red solid line) were presented for anterior PCun, LIPL, LIFG, and LMFG (from left to right). PCun = precuneus; LIPL = left inferior parietal lobule; LIFG = left inferior frontal gyrus; LMFG = left middle frontal gyrus; UWS = unresponsive wakefulness syndrome; UWS-E = patients who emerged from UWS at least three months after the experiment performance; UWS-R = patients who remained in UWS at least three months after the experiment performance; DC = degree centrality.

**Figure 4.**
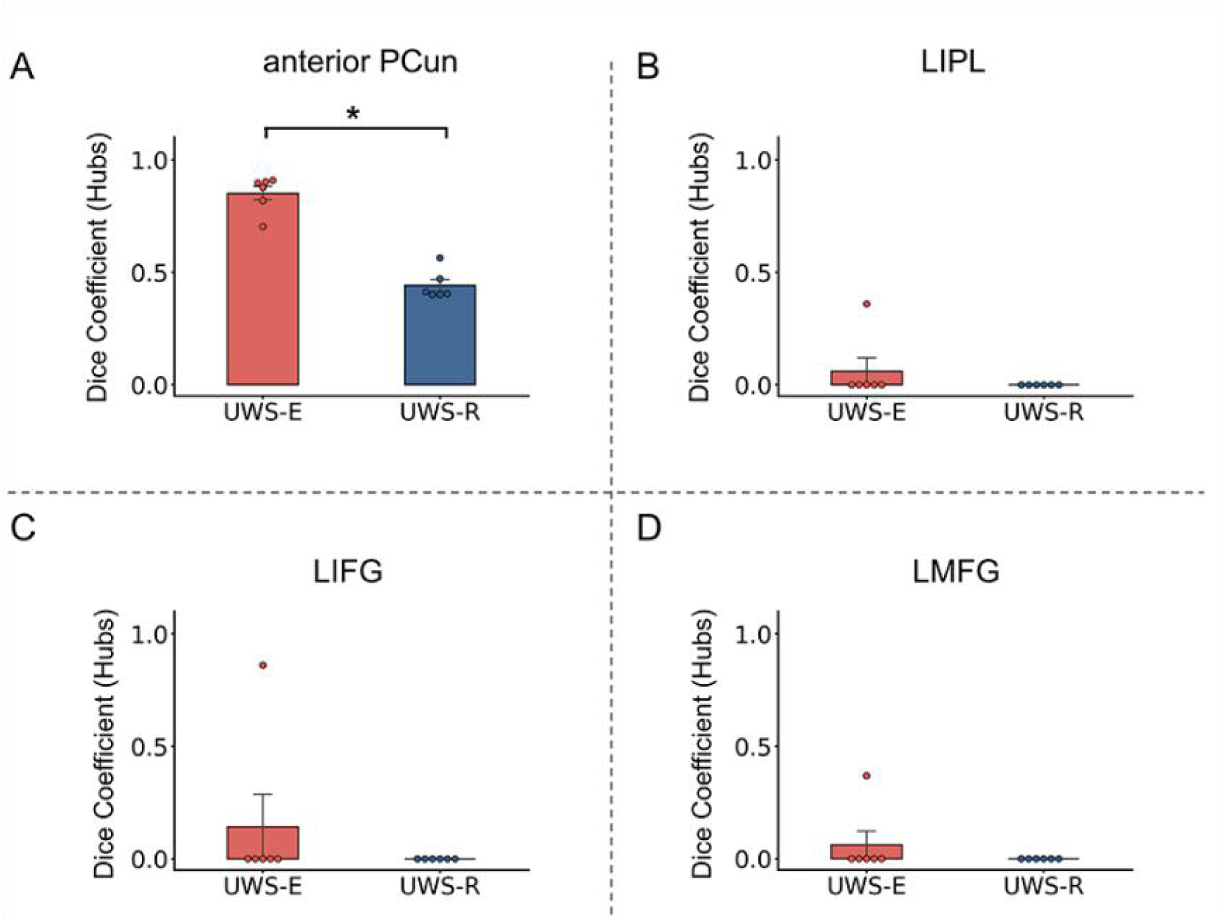
Spatial similarity of cortical hub distribution between the two patient subgroups and the validation groups. Results of spatial similarity measured with dice coefficient was presented within the anterior PCun (A), LIPL (B), LIFG (C), and LMFG (D). PCun = precuneus; LIPL = left inferior parietal lobule; LIFG = left inferior frontal gyrus; LMFG = left middle frontal gyrus; UWS = unresponsive wakefulness syndrome; UWS-E = patients who emerged from UWS at least three months after the experiment performance; UWS-R = patients who remained in UWS at least three months after the experiment performance; * indicates p < 0.05 FDR corrected.

### Similarity of functional connectivity patterns among UWS-E, deep sleep and anesthesia state

For the connectivity patterns in UWS-E (figure 5B) and all six validation groups (figure 5D-I), widespread brain areas within the sensorimotor network (SMN) were functionally connected with the anterior PCun, which is similar as the healthy controls (figure 5A). Whereas for UWS-R patients, the functional connectivity with the anterior PCun showed a different pattern (figure 5C). This result was further validated with the group-level comparison of the dice coefficient (shown in figure 6), which revealed a significantly higher similarity for the UWS-E than UWS-R with the validation groups (p < 0.05).

**Figure 5.**
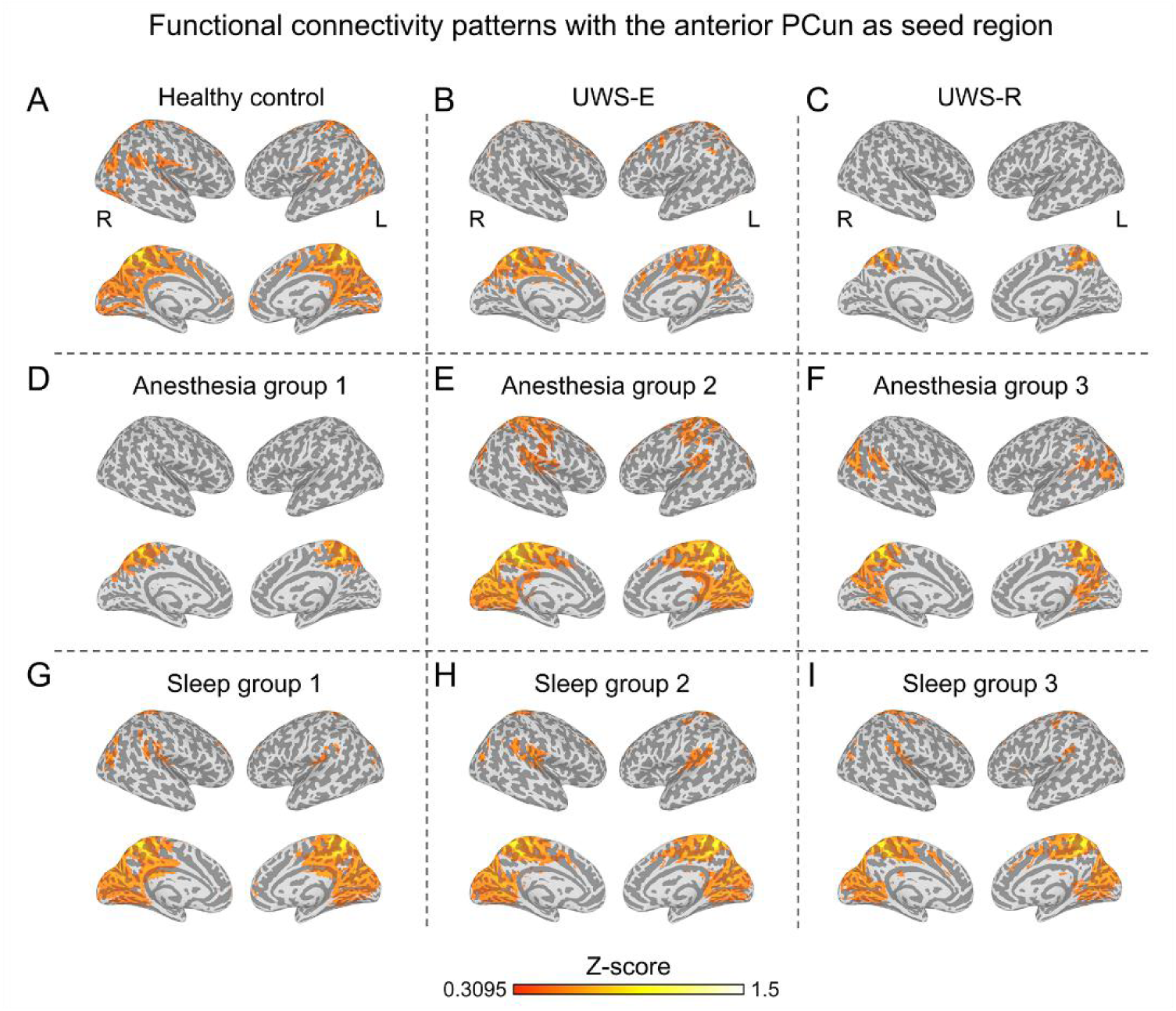
Functional connectivity of the anterior precuneus. Average functional connectivity map with a threshold at a Z value of 0.3095 was displayed for healthy control group (A), UWS-E (B), UWS-R (C), all three anesthesia groups (D-F), and all three deep sleep groups (G-I). UWS = unresponsive wakefulness syndrome; UWS-E = patients who emerged from UWS at least three months after the experiment performance; UWS-R = patients who remained in UWS at least three months after the experiment performance.

**Figure 6.**
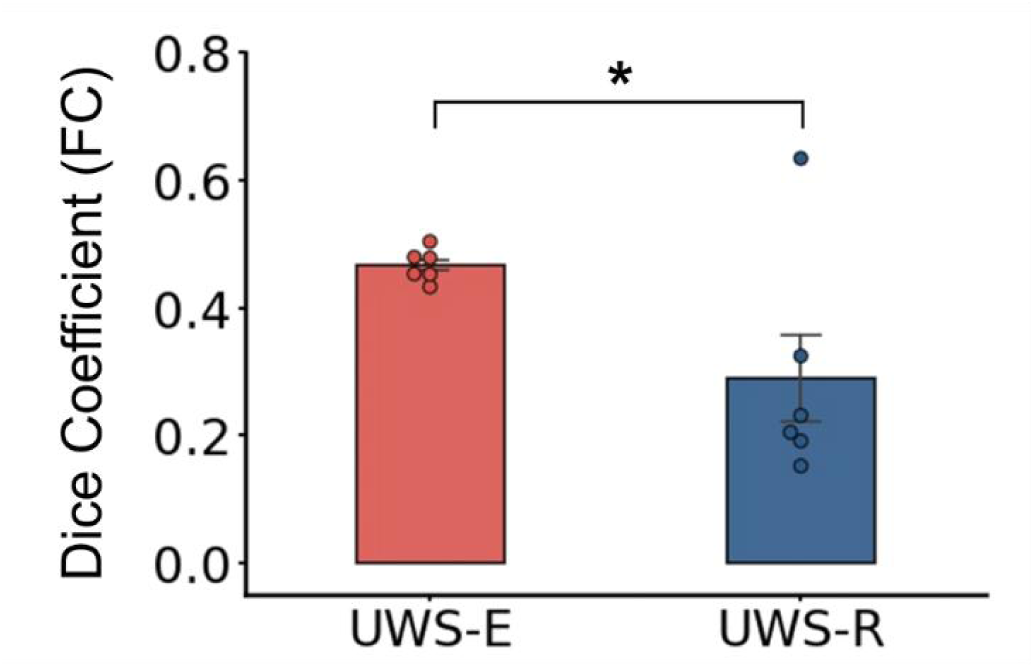
Comparison of the dice coefficient of the functional connectivity with the anterior precuneus between UWS-E and validation groups, as well as UWS-R and validation groups. UWS = unresponsive wakefulness syndrome; UWS-E = patients who emerged from UWS at least three months after the experiment performance; UWS-R = patients who remained in UWS at least three months after the experiment performance. * indicates p < 0.05.

## Discussion

The current study investigates the cortical hubs related to consciousness recovery using patients with UWS. The results showed four cortical hubs including the anterior PCun, LIPL, LIFG and, LMFG exhibited significantly higher degree centrality in UWS-E than UWS-R. This indicated that differential hub characteristics could discriminate recoverable and unrecoverable UWS. Furthermore, among these four hub regions, the anterior PCun consistently showed higher similarity in hub characteristics as well as connectivity patterns, among various recoverable unconscious states, including UWS-E, deep sleep and anesthesia, but not in UWS-R. Given that the cortical hub regions identified in the current study could distinguish the patients in UWS who recover consciousness after 3 months from those who do not, they could serve as potential target sites for UWS treatments using noninvasive stimulation, such as repetitive transcranial magnetic stimulation and transcranial direct-current stimulation.^30^

In the current study, the anterior PCun was found to be a hub that could be related to patients’ recovery from UWS. This is supported by the previous study which showed that the anterior PCun was considered to play an important role in multisensory information integration, which was found associated with bodily awareness.^31^ Furthermore, this region had more functional connections to the SMN in UWS-E than in UWS-R. This finding is supported by previous studies which indicated that the anterior PCun was functionally connected to the SMN, which was regarded as a sensorimotor region.^32,33^ The SMN is mainly responsible for sensorimotor information integration, which has been indicated to play a crucial role in supporting consciousness.^5,6^ In particular, it has been reported that the ability of somatosensory discrimination was found strongly correlated with the UWS patients’ clinical outcomes,^34^ and repetitive transcranial magnetic stimulation^35^ and transcranial direct-current stimulation^36^ to the SMN in patients with DOC could cause clinical improvement.

More interestingly, the finding of similar hub characteristics and functional connectivity patterns for anterior PCun in all recoverable unconscious states (deep sleep and anesthesia) was also supported by previous findings. For instance, precuneus activity was found preserved or increased during sedation^37,38^ and deep sleep state.^39,40^ Our study extended these findings by showing that in conditions where consciousness could be restored (UWS-E, deep sleep, anesthesia), the connectivity patterns with the anterior PCun was highly similar, which were different from UWS-R. Taken together, the current results indicated that the anterior PCun could be a sensorimotor hub that plays a crucial role in preserving the functional integrity that supports the recovery of consciousness in general.

In the current findings, the LIPL also showed higher degree centrality values for UWS-E than for UWS-R. This finding is consistent with previous literature, which showed that the LIPL connectivity was positively correlated with outcomes in DOC patients.^8,41^ As a key node of the default mode network (DMN), the LIPL was indicated to integrate multisensory information from external pathways including visual, auditory, and somatosensory processing.^42^ Combining the current result, it is possible that a preserved ability of external sensory information integration could be necessary for the recovery of consciousness. On the other hand, our results showed extensive hub distributions within the midline components of DMN for both UWS-E and UWS-R as well as deep sleep and anesthesia states, which include the posterior cingulate cortex (PCC), ventral precuneus, and medial prefrontal cortex (MPFC). This is consistent with previous studies which showed that spontaneous synchronized DMN activity was preserved in patients in coma,^9,43^ UWS,^44,45^ deep sleep stage,^39,40^ as well as in the propofol and sevoflurane-induced sedation.^46–48^ This results indicated that each region of DMN might have distinct relationship with recovery of consciousness from others.

Furthermore, LMFG and LIFG were also found to be hub regions in UWS-E. LMFG and LIFG are regions within the frontoparietal network (FPN), and this result agrees with the proposed mesocircuit fronto-parietal model, which proposes that the activation of the FPN is strongly associated with the restoration of function within the anterior forebrain mesocircuit, that could support graded reemergence of behavioral responsiveness across different levels of DOC.^49^ The current finding supported this model in that the preserved functional integration in the frontal cortex was relevant for consciousness recovery in UWS patients.

One issue should be noted, in the deep sleep and anesthesia states, only the anterior PCun exhibited similar hub characteristics, but not the LIPL, LIFG, and LMFG. This could be due to some true difference among the different states, but could also be due to random factors, such as the difference of laboratories. Therefore, future studies might be needed to further confirm the current results.

In conclusion, the current results showed that preserved cortical hub regions were relevant to the clinical outcome in UWS patients, including the anterior PCun, LIPL, LIFG and LMFG. Moreover, only the anterior PCun reliably demonstrated hub characteristics and a high similarity among recoverable unconsciousness states, including UWS-E, deep sleep and anesthesia. Taken together, the current study offered new insight into the neural mechanism of the recovery of consciousness in patients with UWS, and the identified cortical hubs could provide potential treatment target for patients with UWS.

## Author Contributions

All authors contributed to the critical revision and final approval of the manuscript version for publication. PQ, XW, HW, YM and GN designed this study and wrote the manuscript. PQ and HW performed the data analysis. CW, XW, JZ, ST, ZH and SF also contributed to the data acquisition. HW, XW, AGH, SF, ZQ and DZ also contributed to writing of the manuscript.

